# N-acetyl-L-leucine treatment attenuates neuronal cell death and suppresses neuroinflammation after traumatic brain injury in mice

**DOI:** 10.1101/759894

**Authors:** Chinmoy Sarkar, Nivedita Hegdekar, Marta M. Lipinski

**Affiliations:** Shock, Trauma and Anesthesiology Research (STAR) Center, Department of Anesthesiology; Department of Anatomy and Neurobiology, University of Maryland School of Medicine, Baltimore, MD 21201

**Author notes:** Correspondence to MML, and CS.

**Keywords:** autophagy, traumatic brain injury (TBI), neuronal cell death, neuroinflammation, N-acetyl-leucine (NAL), N-acetyl-DL-leucine, N-acetyl-L-leucine, N-acetyl-D-leucine, concussion, IntraBio, intracranial injury

## Abstract

Traumatic brain injury (TBI) is a major cause of mortality and long-term disability around the world. Even mild to moderate TBI can lead to lifelong neurological impairment due to acute and progressive neurodegeneration and neuroinflammation induced by the injury. Thus, the discovery of novel treatments which can be used as early therapeutic interventions following TBI is essential to restrict neuronal cell death and neuroinflammation. In this study, we demonstrate that orally administered treatment with N-acetyl-L-leucine (NALL) led to attenuation of cell death and reduced the expression of neuroinflammatory markers after controlled cortical impact (CCI) induced experimental TBI in mice. Our data indicate that partial restoration of autophagy flux mediated by NALL may account for the positive effect of treatment in the injured mouse brain. Taken together, our study indicates treatment with NALL would be expected to be beneficial in restricting neuronal death and hence improving neurological function after injury, and is a promising novel, neuroprotective drug candidate for the treatment of TBI.

## Introduction

Traumatic brain injury is a major health concern all over the world. Almost 27 million new cases of TBI were reported in 2016 with an incidence rate of 369 per 100,000 populations in the world^1^. Between 1990 and 2016, the prevalence of TBI increased by around 8.4%^1^. In the US around 1.7 million cases of TBI are reported each year^2,3^. People of different ages are affected by brain injuries that lead to lifelong disabilities and premature death due to severe neurodegeneration^2,4,5^. This brings immense emotional distress to the family members of the affected individual and puts a huge financial burden on the society^3^, indicating the need for development effective therapeutic strategies to restrict neuronal loss after TBI.

In TBI, primary mechanical injury to the brain, for example due to a fall or a vehicle accident, triggers a cascade of secondary events that include excitotoxicity, perturbation in calcium homeostasis, oxidative stress and activation of caspases and calpain, leading to acute and progressive neurodegeneration^5^. TBI also induces highly complex inflammatory responses that initiate rapid proliferation of resident microglial cells and infiltration of macrophages into the injured brain^6,7^. These activated immune cells release a variety of neurotoxic proinflammatory cytokines that further contribute to both acute and chronic neurodegeneration^5-7^. While the primary mechanical injury in TBI is instantaneous and cannot be altered, the secondary injury’s extended timeframe represents a therapeutic opportunity for a treatment aimed at restricting restrict neuronal cell death and at suppressing neuroinflammation after TBI.

Recently, N-acetyl-leucine (NAL), an acetylated derivative of leucine has been shown to improve neurological functions in cerebellar ataxia^8,9^. The racemic mixture, N-acetyl-DL-leucine (ADLL) has been used as a medication for the treatment of acute vertigo and vertiginous symptoms in France since 1957; it is orally available and has an excellent safety profile^10^. Studies in a guinea pig model of acute vertigo showed that ADLL can restore the membrane potential of abnormally polarized neurons of the medial vestibular nucleus^11^. It has also been shown to reduce postural imbalance caused by the vestibular lesion following unilateral labyrinthectomy in rats^12^. *In vitro* electrophysiological studies indicated that such a beneficial role of NAL might be mediated through restoration of previously abnormal membrane potential to a normal state, as demonstrated in vestibular nuclei in guinea pig^8,11^. Promising outcome following NAL treatment in improving ataxic symptoms were also observed among Niemann Pick disease patients, a neurodegenerative lysosomal storage disorder caused by mutation in cholesterol transporting NPC 1 and 2 genes^13,14^. These reports suggest that NAL has a neuroprotective function and led us to hypothesize that NAL treatment may be useful in restricting neuronal cell death after TBI.

In this study we assessed whether NAL was effective at preventing neurodegeneration and neuroinflammation after experimental TBI in mice. Studies on the pharmacological^12^, as well as pharmacokinetic^15^ differences between the enantiomers of NAL have suggested the superior benefit of the isolated L-enantiomer versus the racemate of L and D-enantiomer. Therefore, our investigation was based on the L-enantiomer N-acetyl-L-leucine (NALL).

Our data demonstrate that treatment with oral NALL can attenuate cell death after TBI in a controlled cortical impact (CCI) mouse model. This is associated with a decrease in several neuroinflammatory markers as well as improvement in autophagy flux. These data suggest NALL may represent a promising novel drug candidate for the treatment of TBI.

## Methods

### Controlled cortical impact

All surgical procedures and animal experiments were performed as per the protocols approved by the Animal Care and Use Committee of the University of Maryland. Controlled cortical impact (CCI) induced TBI was performed in male *C57BL6/J* mice (20-25g)^16^. Briefly. After a 10-mm midline incision was made over the skull, the skin and fascia were retracted, and a 4-mm craniotomy was made on the central aspect of the left parietal bone of mice under surgical anesthesia (2-3% isoflurane evaporated in a gas mixture containing 70% N_2_O and 30% O_2_). Moderate injury was induced in the exposed brain by a custom microprocessor-controlled and compressed air driven pneumatic impactor of 3.5 mm diameter tip with the impact velocity of 6 m/s and a deformation depth of 2 mm.

### NAL treatment

N-Acetyl-L-Leucine (441511, Sigma) was dissolved in ethanol to prepare 50mg/ml solution which was then diluted in water to get 10mg/ml. Around 0.25 ml of NALL solution (10mg/ml) was orally administered in mice at 100mg/kg dose (2.5mg NAL/25g mouse) via oral gavage each day after CCI induced TBI for 4 days. Mice were also fed with NALL at 0.5mg/kg chow powder chow mixed with diet gel for up to 7 days after injury.

### Western Blot Analysis

Around 5 mm tissue of ipsilateral cortex around the site of injury from TBI mice or the corresponding tissue of same volume around the same cortical region from sham mice were dissected and processed as described^17^. Tissue lysates were resolved on 4-20% SDS-PAGE gels (Bio-Rad, 5671095) and transferred to PVDF membrane (Millipore, IPVH00010) using semi-dry transfer (Bio-Rad), blocked with 5% nonfat milk in tris buffered saline with 0.05% tween 20 (TBST), probed with primary antibodies in 1% BSA in TBST overnight at 4 °C and incubated with HRP-conjugated secondary antibodies (KPL, 474–1506, 474–1806, 14– 16–06 and 14–13–06) in blocking solution at room temperature for 1h. Protein bands were detected using chemiluminiscence kit (Pierce, 34076) and visualized using Chemi-doc system (Bio-Rad). Band intensity was analyzed using Image Lab software (Bio-Rad) and normalized to loading control (β-Actin).

Primary antibodies: LC3 (1:1000; Novus, NB100-2220), p62/SQSTM1 (1:1000; BD Bioscience, 610832), β-actin/ACTB (1: 10,000; A1978) and fodrin/spectrin (1:5000; Enzo Life Science International, BML-FG6090).

### TUNEL assay

20 μm frozen sections were cut from vehicle or NAL fed sham and TBI mouse brains fixed with 4% paraformaldehyde (PFA, pH 7.4) and protected in 30% sucrose. TUNEL assay was performed on those frozen brain sections using ApopTag In Situ Apoptosis Detection Kit (Millipore, S7165) as per the manufacturer’s protocol. Images following TUNEL assay were acquired using a fluorescent Nikon Ti-E inverted microscope, at 20X (CFI Plan APO VC 20X NA 0.75 WD 1 mm).

### Real Time PCR

Total RNA isolated from ipsilateral cortex using miRNeasy Mini Kit (Qiagen, Cat No. **217004**) was converted into cDNA using High Capacity RNA to cDNA kit (Applied Biosystem, Cat. No. 4387406) as per manufacturer’s instruction. cDNA TaqMan® Universal Master Mix II (Applied Biosystems, Cat. No. 4440040) was used to perform quantitative real-time PCR amplification as described previously^17^ using 20 × TaqMan® Gene Expression Assay (Applied Biosystems) for the following mouse genes: *Gapdh* (Mm99999915_g1), *NLRP3* (Mm00840904_m1), *NOX2* (Mm01287743_m1), *iNOS* (Mm00440502_m1), *TNFα* (Mm00443258_m1), *IFnβ1* (Mm00439552_s1), *IL-10* (Mm01288386_m1), *IL1b* (Mm00434228_m1), *Arg1* (Mm00475988_m1), *SOCS3* (Mm00545913_s1), *YM-1* (Mm00657889_mH) and *IL-4ra* (Mm01275139_m1) (Applied Biosystems). Reactions were amplified and quantified by using a 7900HT Fast Real-Time PCR System and corresponding software (Applied Biosystems). Relative gene expression normalized to *Gapdh* was calculated based on the comparative Ct method^18^.

### Statistics

All data are presented as mean ± standard error of the mean (SEM). One-way ANOVA was performed followed by appropriate post-hoc test (Tukey’s) as specified in the figure legends. For data with only two groups 2-tailed student t-test with equal variance was used. A *p* value ≤ 0.05 was considered statistically significant.

## Results

### NALL treatment does not affect food intake and body weight of mice after TBI

To investigate if NALL treatment can prevent cortical cell death after TBI, we orally fed C57/BL6 mice with NALL for 1 day – 7day following controlled cortical impact induced experimental TBI as depicted in Figure 1A. No significant difference in food intake was observed between the sham and TBI mice fed with NALL (Fig. 1B). While a slight decrease in body weight was detected in vehicle treated TBI mice, no appreciable change in body weight was observed in NALL treated TBI mice (Fig. 1C). These results clearly suggest that inclusion of NALL in chow does not negatively affect food intake or body weight in mice with or without TBI.

**Figure 1.**
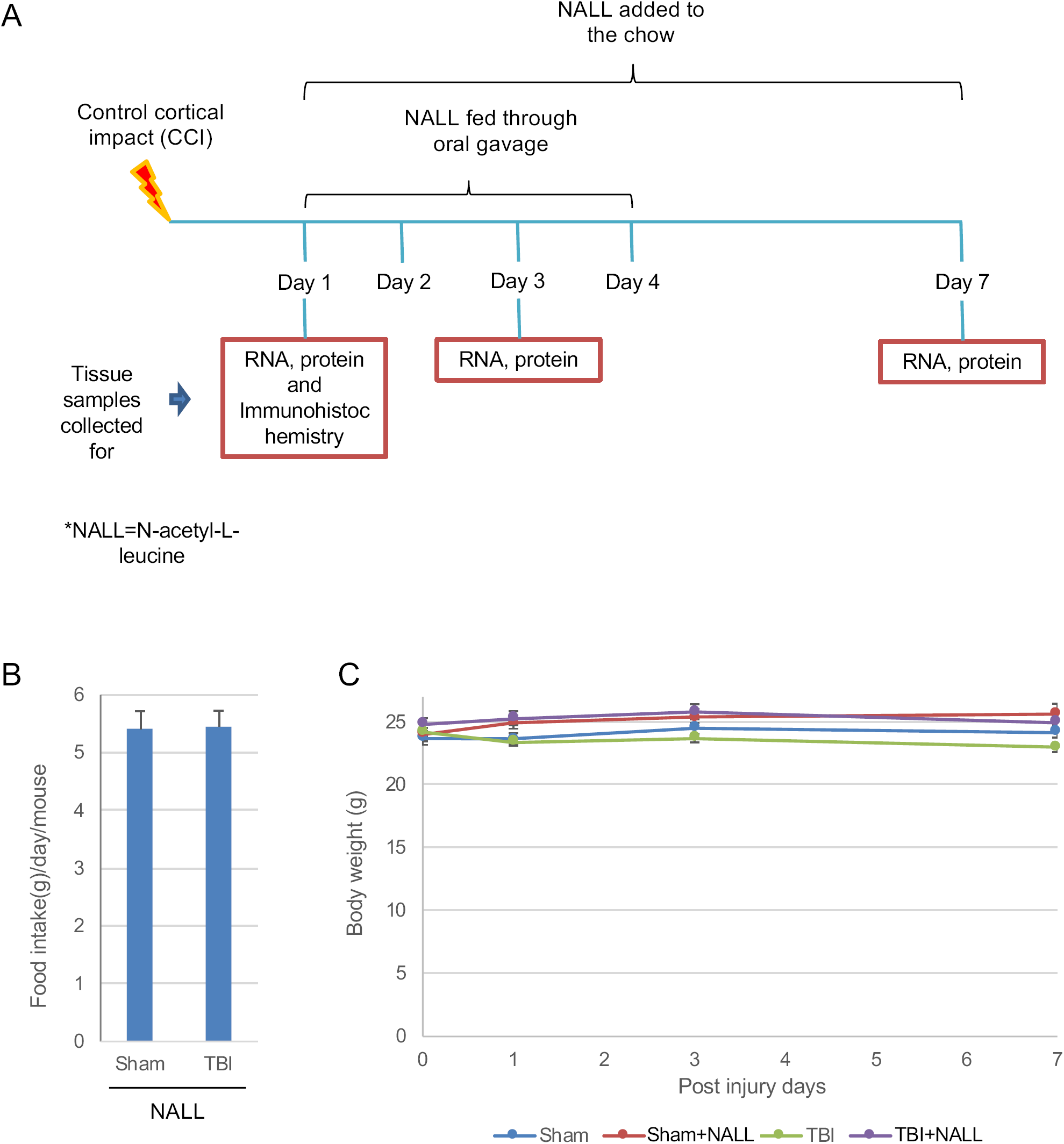
NALL treatment does not affect food intake and body weight in mice. **(A)** NALL treatment strategy. Mice were orally fed with NALL and cortical tissues were collected at days 1, 3 and 7 after TBI for biochemical analyses. **(B)** Amount of NALL containing chow eaten by the mice. No significant changes in food intake were observed in mice fed with NALL containing chow. **(C)** Body weight of sham and injured mice fed with vehicle or NALL. Data are presented as mean ± SEM. n= 7.

### NALL treatment attenuates cortical cell death after TBI

Next, we assessed whether NALL treatment can decrease cortical cell death after TBI. The level of *α*-fodrin cleavage products in NALL, vehicle treated sham, and TBI mouse cortices were determined by Western blot. TBI induces both calpain and caspase mediated cleavage of *α*-fodrin generating 145-150 kDa and 150-120 kDa fragments, respectively^17^. We observed significantly lower level of 145-150 kDa fragments of *α*-fodrin in the NALL fed mouse cortices as compared to the cortices of mice fed with vehicle after TBI, indicating the protective benefit of NALL in preventing cortical cell death (Fig 2A and B). We also detected lower levels of TUNEL positive cells in the brain sections of injured mice fed with NALL as compared to vehicle fed TBI mice (Fig 2C and D). This clearly demonstrated the protective role of NALL in attenuating cell death after TBI.

**Figure 2.**
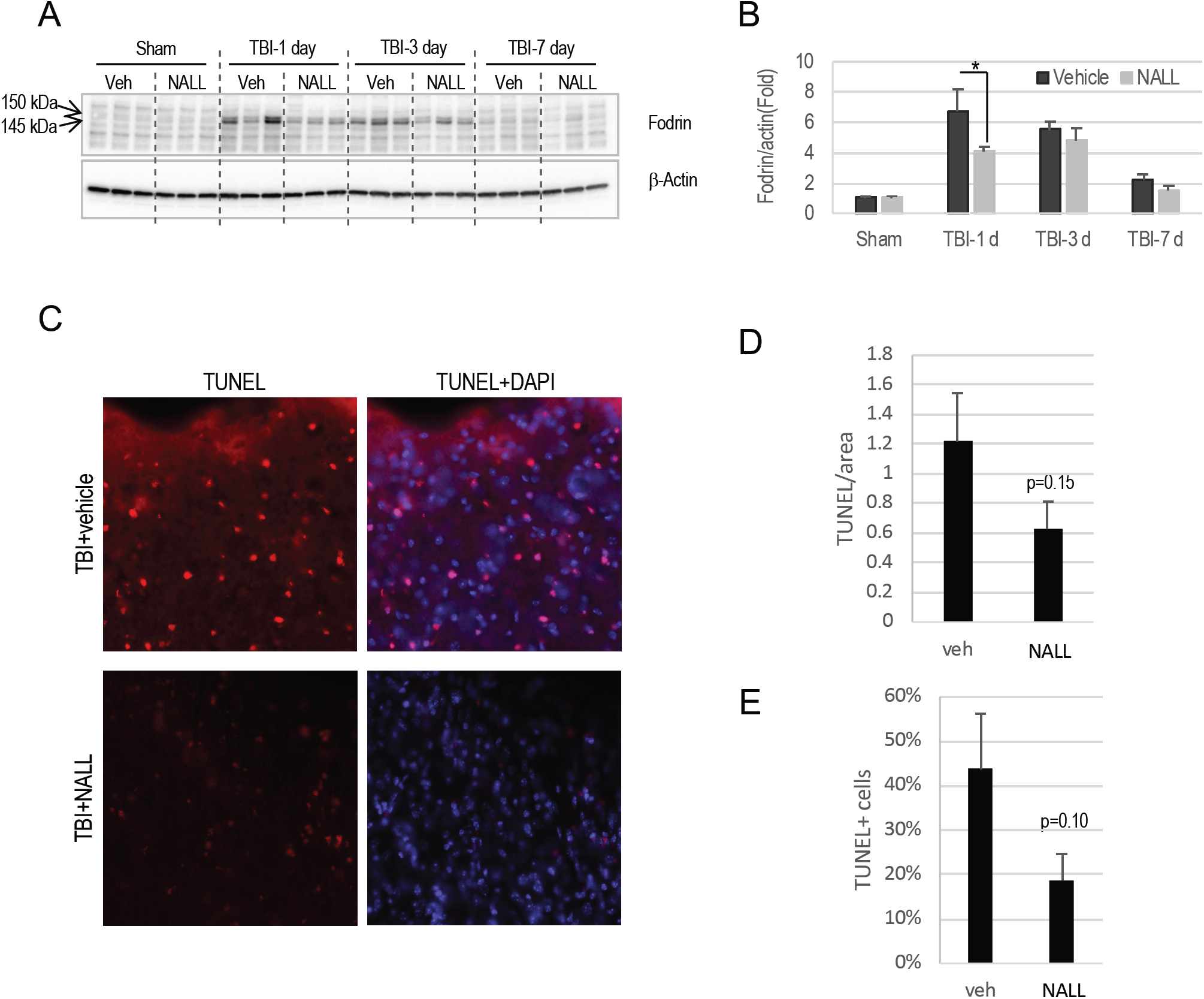
NAL treatment attenuates cell death after TBI. **(A)** Western blot of cortical tissue lysate of sham and TBI mice fed with NALL or vehicle to detect *α*-fodrin breakdown products. **(B)** Corresponding densitometric analysis cleaved bands of *α*-fodrin with respect to actin. Data are presented as mean ± SEM. n=5, **p<0.01 (Two-way ANOVA with Bonferroni posttests). **(C)** 20 × images of vehicle or NALL fed TBI mouse cortical brain sections stained for TUNEL and (D-E) corresponding quantification. Data are presented as mean ± SEM. n=3 for vehicle fed and 4 for NALL fed TBI mice.

### NALL treatment restores autophagy flux after TBI

Recently, we have demonstrated that autophagy, a lysosome dependent cellular degradation process, is disrupted due to lysosomal damage after TBI^17,19^. Autophagy is an important process for neuronal cells as it removes intracellular protein aggregates and damaged organelles^20-22^. We reported that disruption of autophagy flux after TBI causes accumulation of autophagosomal marker LC3-II and autophagic substrate p62/SQSTM1, and is associated with neuronal death^17^. Hence, we assessed if attenuation of cell death following NALL treatment in TBI mice is mediated through the restoration of autophagy flux. The levels of LC3-II and p62/SQSTM1 in the cortical tissue lysates prepared from vehicle and NAL fed naïve and TBI mice were determined by western blot. We detected a significant decrease in the levels of LC3-II in the brain of NALL fed mice as compared to vehicle treated mice at day 1 after TBI (Fig. 3A-B). We also observed a substantial decrease in p62/SQSTM1 level in the cortex of mice fed with NALL as compared vehicle treated mice after TBI (Fig. 3A and C). This demonstrated that NALL can improve autophagy flux and thereby restore its neuroprotective function in the mouse cortices after TBI.

**Figure 3.**
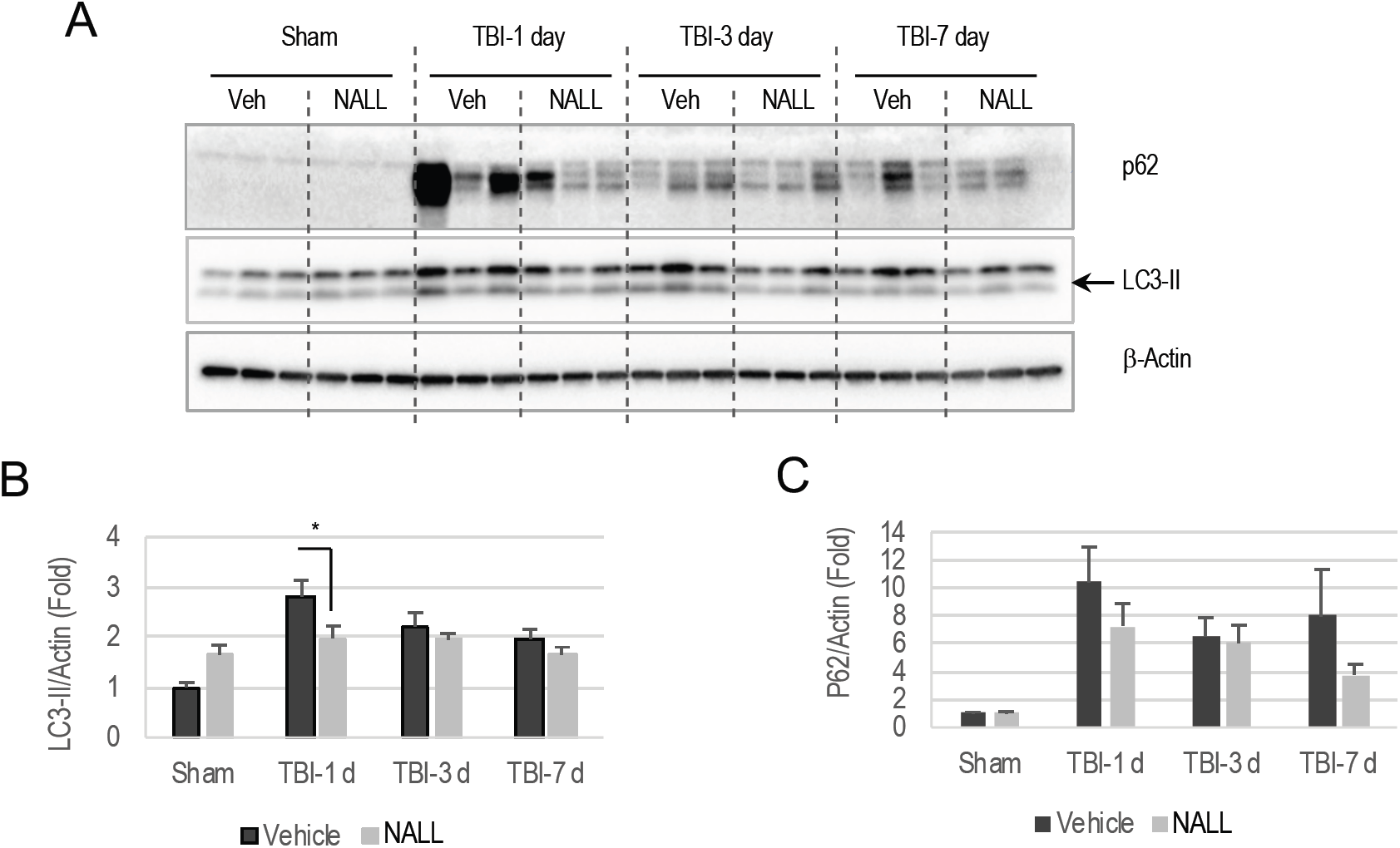
NALL treatment restores autophagy flux after TBI. **(A)** Western blot of cortical tissue lysate of sham and TBI mice fed with NALL or vehicle for autophagosomal marker LC3 and autophagic cargo proteins p62/SQSTM1 and **(B)** corresponding densitometric analysis. Data are presented as mean ± SEM. n=5, *p<0.05 (Two-way ANOVA with Bonferroni posttests).

### NALL reduces inflammatory cytokines in TBI mouse brain

We examined whether NAL treatment can reduce neuroinflammation in the brain following TBI. We determined the expression level of M1-type pro-inflammatory markers Nos2, Nlrp3, IL-1β, TNF, IFNβ and Nox2 in the perilesional area in TBI mouse cortices by real time rt-PCR (Fig. 4A-F). We observed markedly elevated levels of all markers, which started from day 1 and peaked at day 3 after injury in all TBI mouse cortices, irrespective of treatment. However, we detected a significant decrease in the mRNA level of IFNβ and Nox2 in the cortices of NAL fed mice as compared to vehicle fed TBI mouse cortices (Fig 4A-F). Similar to M1-type markers, we observed significantly higher expression of M2-type markers Socs3, YM-1, IL4ra and Arg-1 in the cortical tissue of all TBI mice as compared to sham animals (Fig 5A-D). Among these markers, NAL treatment significantly lowered levels of Arg-1 in the injured mouse cortex as compared to vehicle fed TBI mice. However, levels of Socs3, YM-1 and IL-4ra remained unaltered in NAL fed mouse cortices as compared to sham mice. Taken together, these results demonstrate that NAL reduces expression of several inflammatory markers, thus indicating overall decrease in neuroinflammation following TBI in mice.

**Figure 4.**
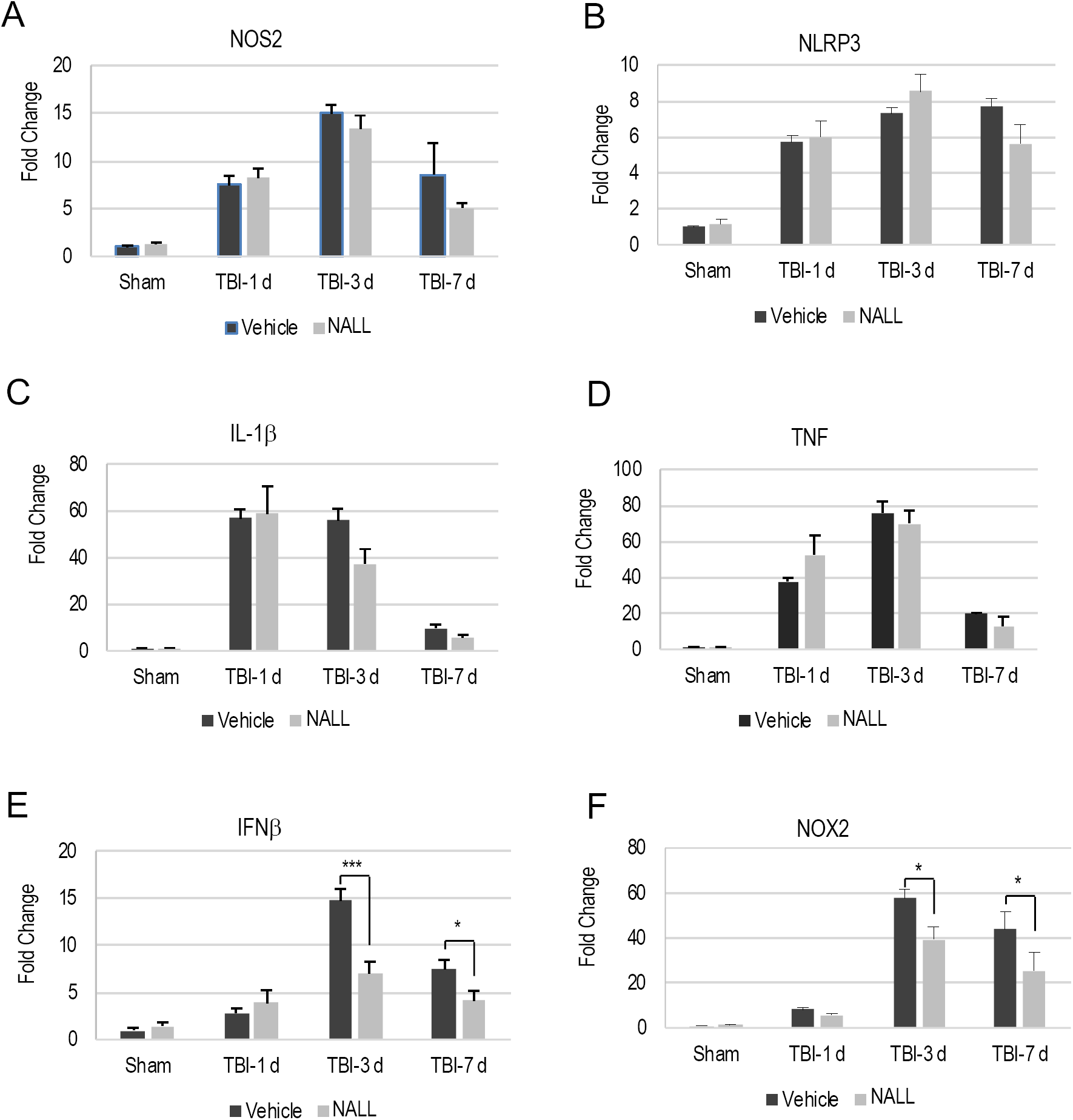
NAL treatment reduces inflammatory (M1) markers in the injured mouse brain. Relative mRNA levels of **(A)** *NOS2*, **(B)** *NLRP3*, **(C)** *IL-1β*, **(D)** *TNF-α*, **(E)** *IFN-β* and **(F)** *NOX2* in the cortices of sham and TBI mice fed with NALL or vehicle. Data are presented as mean ± SEM. n=5, ***p<0.001, *p<0.05 (Two-way ANOVA with Bonferroni posttests).

**Figure 5.**
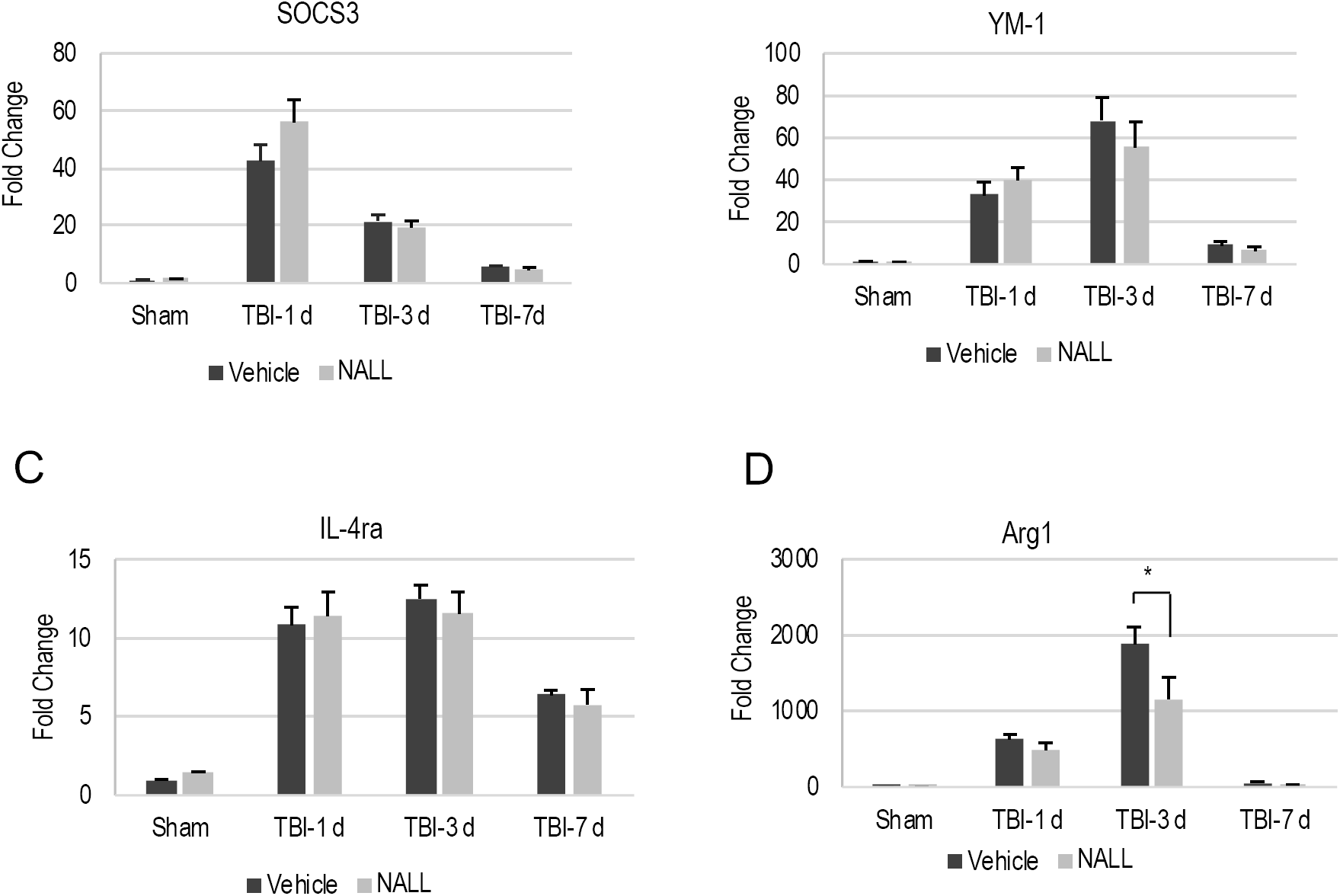
NAL treatment reduces inflammatory (M2) markers in the injured mouse brain. Relative mRNA levels of **(A)** *SOCS3*, **(B)** *YM-1*, **(C)** *IL-4ra* and **(D)** *Arg-1* in the cortices of sham and TBI mice fed with NALL or vehicle. Data are presented as mean ± SEM. n=5, *p<0.05 (Two-way ANOVA with Bonferroni posttests).s

## Discussion

Repurposing of drugs that are already in clinical use for treatment of certain medical conditions can be an effective and rapid way to develop useful therapeutic strategies for the treatment of highly devastating and intractable conditions like TBI. As described, the racemic mixture of N-acetyl-leucine (DL-NAL) has been used orally for more than 60 years in France to treat acute vertigo and dizziness^8^.

It has previously been observed that the L-enantiomer (NALL) is the pharmacologically active enantiomer of the racemate^12^. In addition, differences in the pharmacokinetics of N-acetyl-DL-leucine enantiomers indicate chronic administration of the racemate could have negative effects and support the isolated use of N-acetyl-L-leucine^15^. Accordingly, in this study, we investigated NALL and demonstrated its neuroprotective function in the mouse model of experimental TBI.

Early inhibition of cell death is very important in managing the devastating neurological outcome following TBI. Marked attenuation of cell death was observed during the acute phase at day 1 after TBI in mice that were treated orally with NALL. Our data demonstrate that this might be mediated through the activation or restoration of autophagy flux by NALL after TBI. Autophagy has a neuroprotective function^23^. Thus, early restoration of autophagy flux after TBI by NALL treatment would be expected to be therapeutically extremely beneficial in restricting progressive neuronal death and hence in improving neurological function after TBI.

NALL treatment also reduced neuroinflammation in the injured cortices of TBI mice. Acute and prolonged neuroinflammation contribute to neuronal death after TBI. Thus, attenuation of neuroinflammation by NALL treatment in mice after TBI provides additional neuroprotective function in brain trauma. The observed decrease in Nox2 expression may be particularly significant as its gene product is a known mediator of ROS production and its genetic deletion or pharmacological inhibition is neuroprotective after TBI^25^. Taken together, our study clearly demonstrates the beneficial role of NALL treatment in restricting neuronal loss and in attenuating neuroinflammation after TBI.

## Acknowledgements

This research was supported by IntraBio Inc. The funders had no role in data collection or analysis.

